# QTL mapping: an innovative method for investigating the genetic determinism of yeast-bacteria interactions in wine

**DOI:** 10.1101/2021.01.21.427107

**Authors:** Louise Bartle, Emilien Peltier, Joanna Sundstrom, Krista Sumby, James G Mitchell, Vladimir Jiranek, Philippe Marullo

## Abstract

The two most commonly used wine microorganisms, *Saccharomyces cerevisiae* yeast and *Oenococcus oeni* bacteria, are responsible for completion of alcoholic and malolactic fermentation (MLF), respectively. For successful co-inoculation, *S. cerevisiae* and *O. oeni* must be able to complete fermentation, however, this relies on compatibility between yeast and bacterial strains. For the first time, Quantitative Trait Loci (QTL) analysis was used to elucidate whether *S. cerevisiae* genetic makeup can play a role in the ability of *O. oeni* to complete MLF. Assessment of 67 progeny of an *S. cerevisiae* SBxGN cross, co-inoculated with a single *O. oeni* strain, SB3, revealed a major QTL linked to MLF completion by *O. oeni*. This QTL encompassed a well-known translocation, XV-t-XVI, that results in increased *SSU1* expression and is functionally linked with numerous phenotypes including lag phase duration and sulfite export and production. A reciprocal hemizygosity assay was performed to elucidate the effect of the gene *SSU1* in the SBxGN background. Our results instead revealed a strong effect of *SSU1* haploinsufficiency on *O. oeni*’s ability to complete malolactic fermentation during co-inoculation, and paves the way for the implementation of QTL mapping projects for deciphering the genetic bases of microbial interactions.

**Key points:** For the first time QTL analysis has been used to study yeast-bacteria interactions.

A QTL encompassing a translocation, XV-t-XVI, was linked to MLF outcomes.

*S. cerevisiae SSU1* haploinsufficiency positively impacted MLF by *O. oeni.*

## Introduction

Fermented beverages are the result of biotransformation of complex organo-chemical matrices by microbial communities of moulds, yeast, bacteria and bacteriophage (Bokulich and Bamforth 2013; Mounier et al. 2008; Renouf et al. 2007). Within these communities, the growth rate and metabolic activities of each microbial species depends on (i) the biochemical composition of the medium, (ii) the physicochemical conditions of the process (e.g. converting sugars to ethanol during juice fermentation), (iii) the physiological state of the microbes and (iv) cell-to-cell contact and metabolite interaction among microbial species.

Fermenting grape juice is a fast-changing environment that is especially interesting for studying how the two most common wine microbes, the yeast *Saccharomyces cerevisiae* and lactic acid bacterium *Oenococcus oeni*, coexist and interact. In an oenological context, microbial interactions may affect the final profile of volatile compounds (Chasseriaud et al. 2018, Renault et al. 2015, 2016), that subsequently can be detected as changes in wine sensorial complexity (Tempère et al. 2018). The importance of microbial interactions in wine is evident from the wide number of studies focusing on co-inoculated or sequential inoculation of *S. cerevisiae* and *O. oeni*, with the aim of decreasing overall fermentation time while maintaining or increasing wine quality (Abrahamse and Bartowsky 2012; Cañas et al. 2012, 2015; Knoll et al. 2012).

The mechanisms of yeast-lactic acid bacteria (LAB) interactions during juice fermentation have been reviewed recently (Bartle et al. 2019). Broadly, microbial interactions may include cell-cell contact (Nissen et al. 2003, 2004; Renault et al. 2013) or production of small metabolites (Renault et al. 2009; Sadoudi et al. 2012) and macromolecules (Comitini et al. 2005; Jarosz et al. 2014) that can inhibit and/or activate the growth and activity of interacting microbes. Understanding the molecular mechanisms of yeast-LAB interactions is a challenging task due to the confounding effects of the evolving complex growth environment and microbial metabolite production, but the benefits of such work include optimisation of yeast-LAB co-inoculation strategies for implementation in wineries.

*S. cerevisiae* and *O. oeni* interactions can affect their ability to complete alcoholic (AF) and malolactic fermentation (MLF), respectively (Bartle et al. 2019). Yeast may produce metabolic compounds that can inhibit LAB growth, including ethanol (Capucho and San Romño 1994; Gao and Fleet 1995; Guzzo et al. 2000), SO_2_ (Osborne and Edwards 2006), short and medium-chain fatty acids (Alexandre et al. 2004; Capucho and San Romño 1994), and antimicrobial peptides (Atanassova et al. 2003; Mendoza et al. 2010; Nehme et al. 2010). Yeast and LAB also have the potential to interact physically in the form of mixed species biofilms (Arroyo-López et al. 2012; Bartle et al. 2019; Furukawa et al. 2010) or through co-aggregation (Furukawa et al. 2011), though to date there have not been reports of this occurring between *S. cerevisiae* and *O. oeni*.

In addition to chemical and physical interactions, *S. cerevisiae* gene expression is affected by co-inoculation with *O. oeni* (Rossouw et al. 2012). Differential gene expression in *S. cerevisiae* in response to co-inoculation with *O. oeni* included up-regulation of genes related to yeast stress response and possible competition for sulfur compounds compared to *S. cerevisiae* alone (Rossouw et al. 2012). Several studies have also reported strain-specific compatibility between yeast and LAB during co-inoculation (Abrahamse and Bartowsky 2012; Antalick et al. 2013; Arnink and Henick-Kling 2005; Comitini and Ciani 2007; Rossouw et al. 2012; Tristezza et al. 2016). Considering the strain specificity of compatibility outcomes, the intraspecific genetic variability of interacting species requires further investigation and analysis. To our knowledge, the identification of genetic variations to explain “strain compatibility” is a novel approach.

For *S. cerevisiae*, the genetic determinism of any complex trait can be investigated by mapping quantitative trait loci (QTLs) in a segregating progeny (Liti and Louis 2012). In the context of wine, this strategy has been used for elucidating the genetic basis of many traits of industrial interest (Peltier et al. 2019) including acetic acid production (Salinas et al. 2012), rate of nitrogen uptake (Brice et al. 2014; Jara et al. 2014), resistance to stuck fermentation (Marullo et al. 2019), resistance to low pH (Martí-Raga et al. 2017), and the production of aroma compounds (Eder et al. 2018; Huang et al. 2014; Roncoroni et al. 2011; Steyer et al. 2012). To date, QTL mapping has been performed for single pure cultures focusing on traits related to yeast fitness or effect on wine quality. However, this strategy may be applied to any trait resulting in measurable phenotypic variability. In the present work, we applied QTL mapping to delineate how *S. cerevisiae* genetic variability may affect the success of MLF in co-inoculated fermentations with a commercial strain of *O. oeni*.

## Materials and methods

### Media

Shiraz juice: Shiraz grapes (2017 vintage, Coombe vineyard, The University of Adelaide, Waite Campus, Urrbrae, South Australia) were harvested, de-stemmed, crushed and left to macerate at 0 °C for 7 days to enable polyphenolic extraction. Shiraz must be pressed and the juice stored at −20 °C until required. No SO_2_ or antibacterial agents were added to the juice during pressing. Prior to experimentation Shiraz juice was filtered (0.45 µm, catalogue # FHT45, Air-Met Scientific, Victoria, Australia) to remove grape matter and solids. Initial measurements of total sugar were estimated by refractometry and sugar reduced to 250 g L^-1^ by addition of water. L-malic acid was increased to 2.5 g L^-1^ by addition of pure L-malic acid and pH was decreased to 3.5 by addition of tartaric acid, followed by addition of 100 mg L^-1^ diammonium phosphate. Finally, the juice was filter sterilised (0.2 µm).

Liquid de Man, Rogosa and Sharpe medium (MRS; catalogue # AM103, Amyl Media, Victoria, Australia), supplemented with 20% apple juice (MRSAJ) was used for growing bacteria prior to inoculation. MRS was autoclaved and sterile apple juice (0.2 µm filtered) added post sterilisation and before use. MRSAJ supplemented with agar (2%) and cycloheximide (0.5%) following sterilisation of the medium, was used for enumeration of bacteria.

All yeast strains were initially streaked for single colonies on YPD agar (2% glucose, 2% peptone, 1% yeast extract, 2% agar) and grown at 28 °C, before growth of single isolates in YPD (2% glucose, 2% peptone, 1% yeast extract) at 28 °C overnight. If required, Geneticin (G418, 100 µg mL^-1^; catalogue # G8168, Sigma-Aldrich, New South Wales, Australia) was added to YPD cultures to select for strains carrying the *KanMX* deletion cassette.

### Strains and fermentations

#### Yeast strains

Strains used in this work are listed in Table 1. QTL analysis was performed using the SBxGN yeast background. SBxGN is the F1-hybrid of SB and GN strains, two diploid, fully homozygous strains derived from the wine starters Actiflore® BO213 and Zymaflore® VL1, respectively (Peltier et al. 2018b). The population used for QTL mapping consisted of 67 haploid progeny clones derived from the hybrid BN, an isogenic variant of SBxGN (Marullo et al. 2007a). These haploid meiotic progenies have been previously genotyped by whole genome sequencing (Martí-Raga et al. 2017). The effect of the gene *SSU1* was assayed using the reciprocal hemizygosity assay by deleting each parental copy of *SSU1* individually in the SBxGN F1-hybrid (Steinmetz et al. 2002a). The reciprocal hemizygous hybrids SΔG092 and GΔS092 were previously obtained as described by Zimmer and colleagues (2014).

**Table 1:** Yeast strains used in this study

| Strain | Comment | Genotype | Origin | Culture collection:<br>WDCM-791 (CRB<br>ISVV, Bordeaux) |
| --- | --- | --- | --- | --- |
| SB | Monosporic clone of<br>Actiflore® BO213<br>(Laffort®, Bordeaux,<br>France) | *HO/HO*, diploid | Peltier et al.<br>(2018b) | CRBO L2001 |
| GN | Monosporic clone of<br>Zymaflore® VL1<br>(Laffort®, Bordeaux,<br>France) | *HO/HO*, diploid | Peltier et al.<br>(2018b) | CRBO L2002 |
| SBxGN | F1 hybrid SBxGN | *HO/HO*, diploid | Peltier et al.<br>(2018b) | CRBO L2003 |
| BN | F1 hybrid hoSBxGN | *HO/ho::kanMx4*, diploid | Marullo et<br>al. (2007b) | CRBO L2004 |
| pop BN | 67 progeny clones of<br>BN (hoSBxGN).<br>Labelled with prefix<br>“CM” followed by an<br>ID number | *ho::kanMx4*, haploids | Marullo et<br>al. (2007b) |  |
| SΔG092 | Hemizygous hybrid<br>isogenic to SBxGN | *ho/ho*, *YPL092<sup>SB</sup>::kanMX4/*<br>*YPL092<sup>GN</sup>*, diploid | Zimmer et<br>al. (2014) | CRBO L2006 |
| GΔS092 | Hemizygous hybrid<br>isogenic to SBxGN | *ho/ho*, *YPL092<sup>GN</sup>::kanMX4/*<br>*YPL092<sup>SB</sup>*, diploid | Zimmer et<br>al. (2014) | CRBO L2007 |

#### Yeast cell concentration and viability

Yeast live cell concentrations were determined by flow cytometry. Yeast were diluted 100-times in sterile phosphate buffered saline (PBS), then stained with propidium iodide at a final concentration of 0.1 mg mL^-1^. Samples were analysed using a Guava® easyCyte™ 12HT flow cytometer (Luminex, Yokohama, Japan; previously Millipore, Darmstadt, Germany). Yeast concentrations were adjusted to inoculate sterile Shiraz juice at a final rate of 5 × 10^6^ live cells mL^-1^.

#### Bacteria

Freeze-dried SB3 (Laffort®, South Australia, Australia) was grown anaerobically in MRSAJ for four days at 30 °C in 20% CO_2_. Twenty-four hours prior to inoculation, bacteria were centrifuged (2,236 x *g*), the supernatant discarded and the cell pellet washed in sterile Shiraz juice before overnight incubation in fresh sterile Shiraz juice at 30 °C. Bacteria were adjusted to an OD_600_ of 0.55 immediately prior to inoculation. For QTL library fermentations, 200 µL of bacterial culture were added to each fermentation vessel manually through a silicone septum with a 21-gauge needle. For the hemizygote fermentations, 200 µL of bacterial culture were transferred from a 96-well deep well plate to each fermentation vessel using an automated fermentation system (developed after performing the QTL experiment).

#### Fermentations

Fermentations were conducted using an automated fermentation system built on an EVO Freedom workdeck (Tecan, Männedorf, Switzerland; Fig. S1: Online resource 2). The system enabled 384 concurrent fermentations at a volume of up to 25 mL. Full details of the system were described by Hranilovic et al. (2018) and can also be found on the University of Adelaide Biotechnology and Fermentation Facility website (https://sciences.adelaide.edu.au/agriculture-food-wine/research/biotechnology-and-fermentation-facility).

Fermentation vessels were filled with 20 mL of sterile Shiraz juice and inoculated with yeast (5 × 10^6^ live cells mL^-1^) followed by LAB inoculation 24 hours later. Sampling occurred daily, and fermentations were homogenised by stirring prior to sampling. For the QTL mapping experiment, both parental strains (SB and GN), the hybrid BN and the 67 haploid progenies were fermented as pure cultures (in duplicate) or co-inoculated with SB3 (in triplicate). To test the effect of *SSU1*, hemizygous and wild type F1-hybrids were assessed in triplicate for both pure and co-inoculated fermentations with SB3.

### Fermentation monitoring

#### Glucose and fructose consumption

Glucose and fructose concentrations were determined enzymatically using commercially available kits (catalogue # K-FRUGL, Megazyme, Bray, Ireland) following methods modified by Walker et al. (2014). Glucose and fructose consumption were used as determinants for progress of alcoholic fermentations, which were deemed complete when total glucose plus fructose concentration was < 4 g L^-1^.

The amount of glucose/fructose consumed over time was modelled by local polynomial regression fitting with the R-loess function, setting the span parameter to 0.8. Five kinetics parameters were extracted from the model, which are described in Table 2.

**Table 2:** AF and MLF measures used to perform QTL mapping, BN progeny evaluation or statistical analysis for comparison of hemizygous strains with their corresponding SBxGN hybrids. Abbreviations, if assigned, are shown below:

| AF measures | Abbreviation | MLF measures | Abbreviation |
| --- | --- | --- | --- |
| Time to complete AF (hours) | *tend-AF* | Residual L-malic acid concentration for yeast alone fermentations (g L<sup>-1</sup>) | Not assigned |
| Time to reach equivalent of 35% CO<sub>2</sub> (175.53 g L<sup>-1</sup> total sugar) | t35-AF | Percentage L-malic acid consumption for yeast-LAB co-inoculation fermentations | *Pct_malic_co* |
| Time to reach equivalent of 50% CO<sub>2</sub> (143.62 g L<sup>-1</sup> total sugar) | t50-AF | Estimated overall L-malic acid consumed by LAB (g L<sup>-1</sup>) | *Malic_acid_LAB_consumed* |
| Time to reach equivalent of 80% CO<sub>2</sub> (79.79 g L<sup>-1</sup> total sugar) | t80-AF | Time to complete MLF (hours) | *tend-MLF* |
| Slope between t50-AF and t80-AF | s50-80-AF |  |  |
| Percentage L-malic acid consumption or production by yeast alone | *Pct_malic_AF* |  |  |

#### L-malic acid concentration

L-malic acid was measured using an enzymatic test kit (catalogue # 4A165, Vintessential Laboratories, Victoria, Australia) with modifications so that a plate-reader/spectrophotometer (Infinite 200 PRO, Tecan, Männedorf, Switzerland) could be used to measure absorbance. Specifically, each well of a 96 well micro-titre plate was dosed with 70 μL buffer (0.1M gly-gly, 0.1M L-glutamate, pH 10), 14 μL nicotinamide adenine dinucleotide (40 mg mL^-1^), 70 μL distilled water, 0.7 μL glutamate oxaloacetate transaminase (800 U mL^-1^) and 5 μL of sample or one of the L-malic acid standards (ranging from 0 − 3.0 g L^-1^). The plate was incubated at 22 °C for 3 minutes and the first absorbance was read at 340 nm; 7 μL of the 1:10 diluted L-malate dehydrogenase (12,000 U mL^-1^) was added and mixed into each well; the plate was incubated at 22 °C for 15 minutes before the second absorbance was measured at 340 nm. L-malic acid in each sample was calculated from standard curves prepared with known L-malic acid concentrations. L-malic acid degradation was used as the determinant for MLF progress. L-malic acid consumed over time was modelled by local polynomial regression fitting with the R-loess function setting the span parameter to 0.75. MLF was deemed complete when L-malic acid concentration was < 0.1 g L^-1^ and designated *tend-MLF* (Table 2).

L-malic acid end point parameters were determined for yeast alone and yeast-SB3 co-inoculation fermentations. These parameters were: percentage of L-malic acid consumed or produced by yeas alone in relation to the starting L-malic acid concentration of 2.5 g L^-1^ (*Pct_malic_AF*), and percentage of L-malic acid consumed by yeast and LAB in co-inoculated fermentations (*Pct_malic_co*).

To estimate the overall L-malic acid reduction by LAB when co-inoculated with yeast, the average concentration of L-malic acid for co-inoculated fermentations at the end of experimentation was subtracted from the average L-malic acid concentration for corresponding yeast-alone fermentations. This parameter was designated *Malic_acid_LAB_consumed*. The end of the experiment was defined as either reduction of L-malic acid to < 0.1 g L^-1^ or approximately 11 days after the first yeast-LAB pair completed MLF.

A summary of all parameters assessed in this study can be found in Table 2.

### Statistical analysis

All statistical analyses were performed using R versions 3.4.4 or higher (R Core Team 2017). Kendall correlation coefficient test was performed using R/stats package v3.6.2 (R Core Team 2017). The QTL mapping analysis was performed with the R/qtl package (Broman et al. 2003) by using the Haley-Knott regression model that provides a fast approximation of standard interval mapping (Haley and Knott 1992). A threshold corresponding to a 5% and 10% false discovery rate (FDR) was computed by performing 1000 permutations in order to assess the significance of the LOD score for QTL peaks (Churchill and Doerge 1994). The overall procedure was described by Peltier et al. (2018b) for multiple environments mapping.

Linear modelling was performed to evaluate the effect of allele, yeast background and translocation on MLF and AF parameters using the following formula:

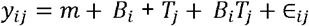

Where *y*_*ij*_ is the value for the background *i* (*i* = 1,2) with translocation *j* (*j* = 1,2), *m* is the overall mean,*B*i is the yeast background effect, *T*_*j*_ is the translocation effect, *B*_*i*_*T*_*j*_ is the interaction effect between yeast background and translocation, ∈ _*ij*_ is the residual error. Tukey post-hoc test (α = 0.05) was used to elucidate differences between ANOVA test groups.

## Results

### Biometric assessment of MLF completion in the SBxGN progeny population

To identify QTLs influencing the completion of MLF, L-malic acid consumption by *O. oeni* was measured in co-inoculated Shiraz grape juice fermentations. Alone, *S. cerevisiae* strains from the BN progeny were able to consume a fraction of L-malic acid (Fig. 1a). The concentration of residual L-malic acid at the end of AF in yeast-alone fermentations ranged from 1.41 g L^-1^ to 2.75 g L^-1^, which corresponded to between 44% consumption and 10% production of L-malic acid in respect to the starting concentration of 2.5 g L^-1^ (Fig. 1b; Table S1: Online resource 1).

**Fig. 1.**
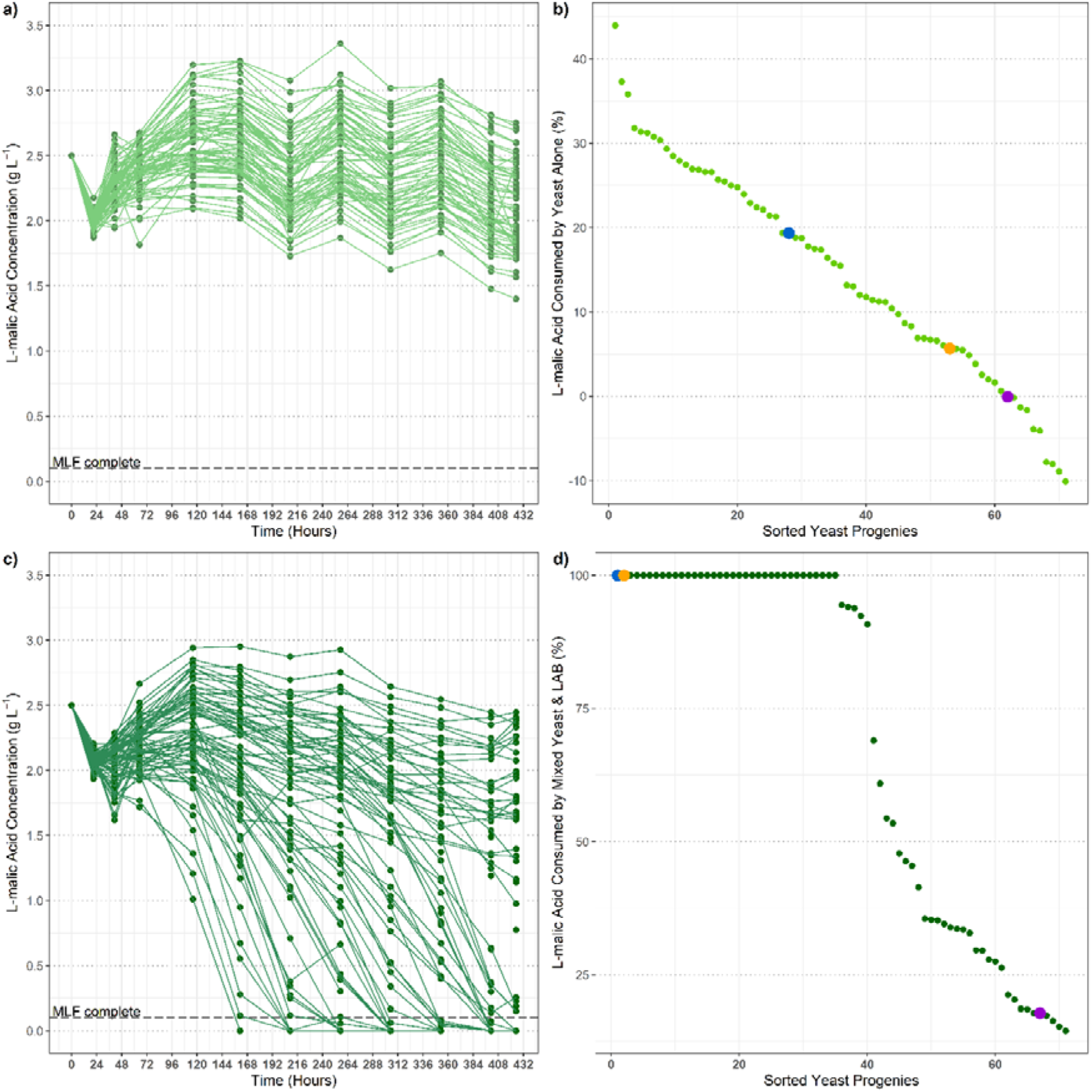
**a)** L-malic acid concentration measured over the course of the experiment for yeast-alone fermentations for the population of 67 SBxGN yeast progeny. Values are the mean of duplicates. **b)** All yeast-alone strains ranked by percentage of L-malic acid consumption (positive %) or production (negative %), measured at the end of the experiment in relation to the starting L-malic acid concentration of 2.5 g L^-1^. Percentages were calculated from the mean of duplicates. Colours indicate yeast parental strains: BN (orange), SB (blue) and GN (purple). All other yeast progeny are shown in green. **c)** MLF progress measured for yeast co-inoculated with SB3 LAB. Values are the mean of triplicates. The horizontal line at 0.1 g L^-1^ indicates when MLF was deemed complete. **d)** All yeast strains in co-inoculations with SB3 LAB were ranked by the percentage of L-malic acid consumed, measured at the end of the experiment. Percentages were calculated from the mean of triplicates. Colours are the same as panel b

In the present study, the focus was the impact of yeast genotype on LAB MLF efficiency. Therefore, we measured L-malic acid consumption over time for fermentations co-inoculated with *S. cerevisiae* strains and LAB SB3 (Fig. 1c). As expected, L-malic acid consumption was much higher for many of the yeast-LAB co-inoculated fermentations. However, SB3 was only able to complete MLF in 39 of the 71 co-inoculated fermentations (Fig. 1d). Since LAB were only able to complete MLF when co-inoculated with some of the SBxGN progenies, this provided evidence of strong yeast-LAB interactions.

Though there were differences in residual L-malic acid across fermentations with different yeast strains, the ability of yeast to consume L-malic acid (as seen for yeast-alone fermentations, Fig. 1a) did not seem to impact MLF completion time by SB3 in co-inoculations. Kendall rank correlation coefficient revealed only a weak positive correlation (0.21, *p* = 0.009) between the amount of L-malic acid consumed by yeast and SB3 MLF completion time.

### QTL mapping

To determine the concentration of L-malic acid consumed only by LAB at the end of the experiment (as defined in materials and methods), the average final L-malic acid concentration found in yeast-LAB co-inoculations was subtracted from the corresponding L-malic acid concentration in yeast-alone fermentations. This new parameter, the L-malic acid consumed by LAB (*Malic_acid_LAB_consumed*), provides a proxy for SB3 MLF efficiency after co-inoculation with different yeast strains. This new parameter has a nearly continuous distribution among the SBxGN progeny (Fig. 2a). Genetic regions linked to the variation of this trait were tracked by applying a linkage analysis. Despite the small number of progenies tested, two loci were linked to this phenotype at an FDR of 10% (Fig. 2b). One QTL peak, located on *S. cerevisiae* chromosome XVI, achieved a LOD score of 7.58 which is highly significant with respect to the threshold value of 4.58 that was estimated by 1000 permutations with an FDR of 5%. An additional peak, located on *S. cerevisiae* chromosome XV had a lower LOD score of 4.02. This LOD score reaches the threshold value of 4.00 which corresponds to an FDR of 10%.

**Fig. 2.**
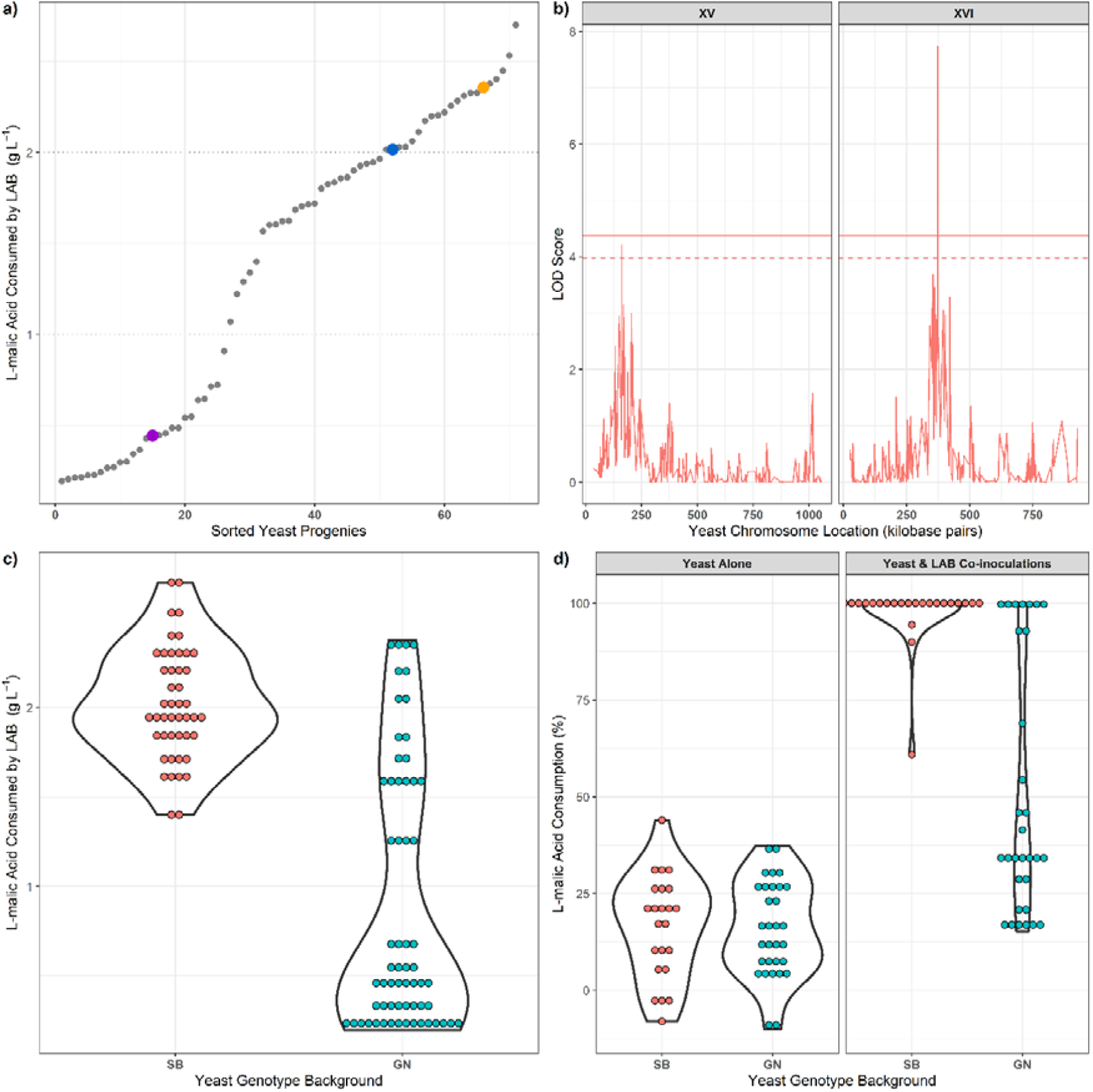
**a)** Yeast ranked by the concentration of L-malic acid that was able to be consumed by SB3 during co-inoculation with each yeast strain. Values determined by subtracting the average residual L-malic acid concentration of the yeast-alone fermentations from the average residual L-malic acid concentration in the corresponding yeast-LAB co-inoculation fermentations. Colours indicate yeast parental strains: BN (orange), SB (blue) and GN (purple). All other yeast progeny are shown in grey. **b)** Genomic location of QTL peaks for the parameter *Malic_acid_LAB_consumed*. Threshold values are estimated from 1000 permutations and 5% FDR, indicated by the solid horizontal line. The dotted horizontal line indicates a LOD score threshold of 4. Significant (peak above threshold) and potential (peaks near a LOD score of 4) QTLs were found on chromosomes XV (left) and XVI (right). **c)** Distribution of yeast progenies with respect to the concentration of L-malic acid consumed by SB3 in co-inoculations with each yeast strain. Progenies are grouped by yeast background (SB, left; GN, right). **d)** Distribution of yeast progenies based on percentage of L-malic acid consumed (measured at the end of experimentation) for yeast alone (left panel) or when co-inoculated with SB3 (right panel). Progenies are grouped by yeast background

The best marker of this linkage analysis was located at genomic position XVI_374156 and was therefore named XVI_374. Due to the density of markers surrounding XVI_374 (6 markers within 817 bp) and *SSU1* spanning the genomic region of chromosome XVI: 373793-375169, there was high confidence in the specificity of *SSU1* being the target of the QTL peak. The inheritance of this marker impacts the consumption of L-malic acid by the LAB SB3. Indeed, co-inoculated fermentations with yeast progenies carrying the SB allele of XVI_374 consumed more L-malic acid than those performed with yeast progenies carrying the GN allele (Fig. 2c). Interestingly, this phenotypic discrepancy is not due to the ability of yeast to consume L-malic acid. Additionally, in yeast-alone fermentations, the inheritance of XVI_374 from either SB or GN did not alter the percentage of L-malic acid consumed by the yeast (Fig. 2d). In contrast, most of the strains containing the yeast SB allele for this QTL allowed SB3 to complete MLF (Fig. 2c). Altogether, these data provide clear evidence that genetic regions of the *S. cerevisiae* genome have a direct impact on the metabolic activity of LAB during co-inoculation.

One peak detected did fall within the 10% FDR (Fig. 2b) and therefore warrants discussion. This peak was located on chromosome XV, at the marker XV_162503 which corresponds to the gene *PHM7*. Although *PHM7* is a gene of unknown function, in the SBxGN background a genetic linkage in this region was predictable since the left arm of chromosome XV is physically linked to chromosome XVI in the strain GN. This well-documented translocation event was previously named XV-t-XVI (Zimmer et al. 2014; Fig. 3) and segregates in a mendelian fashion in the SBxGN progeny. The XV-t-XVI translocation has been demonstrated to impact several fermentation-related phenotypes including lag phase duration, fermentation rate, and SO_2_ production. The molecular basis of such phenotypes is due to modification of the promoter environment of the *SSU1* gene (Fig. 3) that encodes Ssu1p, a transmembrane sulfite efflux pump (Marullo et al. 2020; Peltier et al. 2018b; Zimmer et al. 2014). These previous findings suggest that *SSU1* would be a relevant candidate gene able to explain the yeast-LAB interaction observed in our experiment.

**Fig. 3.**
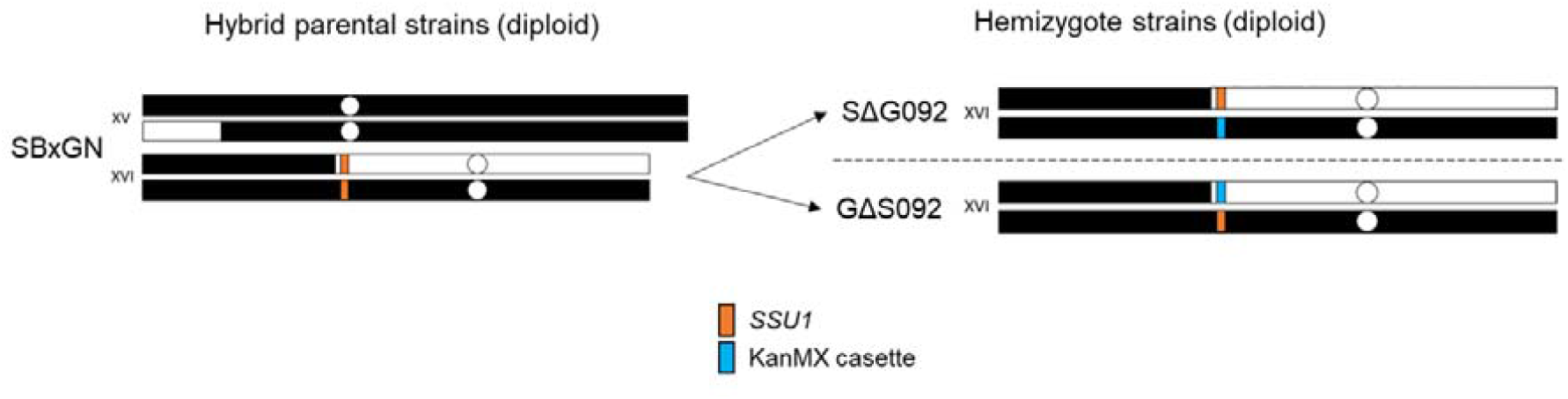
Representation of the translocation located in SBxGN that results in increased *SSU1* gene expression due to reduced proximity between *SSU1* and promotor regions. SBxGN has an XV-t-XVI translocation that leads to a single copy of wild-type XV and XVI chromosomes (all black) and reciprocal XV and XVI translocated chromosomes (black and white). Hemizygous strains SΔG092 and GΔS092 with a single functional *SSU1* allele (orange) were generated to perform a reciprocal hemizygosity assay. Hemizygous strains were created by replacing a single copy of *SSU1* with a KanMX cassette (blue)

### Functional study of a QTL closely related to *SSU1*

To determine the possible influence of *SSU1* on MLF outcome, we applied a reciprocal hemizygosity assay, enabling evaluation of the impact of the two parental alleles in an isogenic context, as proposed by Steinmetz et al. (2002b). To test this hypothesis, we used the hemizygous hybrids SΔG092 and GΔS092 in which one copy of the *SSU1* gene was replaced with the *KanMx4* cassette (Zimmer et al. 2014). Such strains are isogenic to the F1-hybrid SBxGN but carry only one functional copy of *SSU1*: SΔG092 carries the *SSU1* allele of GN while GΔS092 carries the *SSU1* allele of SB. The two hemizygous hybrids were compared to the SBxGN hybrid carrying both functional *SSU1* alleles.

Co-inoculated and yeast-alone fermentations were carried out in the same conditions applied for QTL mapping. In co-inoculated fermentation, all the malolactic fermentations were completed. However, the time taken for SB3 to reduce L-malic acid to < 0.1 g L^-1^ was significantly different depending on the yeast strain (Fig. 4a). Indeed, both hemizygous hybrids enabled shorter MLF completion time (by at least 48 hours) than the parental hybrid SBxGN, revealing a strong haploinsufficiency effect of the *SSU1* deletion. Surprisingly, MLF completion was not significantly impacted by inheritance of either the SB or GN allele (Fig. 4b). This result led to the conclusion that the *SSU1* gene was not the main cause of the SBxGN QTL on chromosome XVI that was based on L-malic acid consumption.

**Fig. 4.**
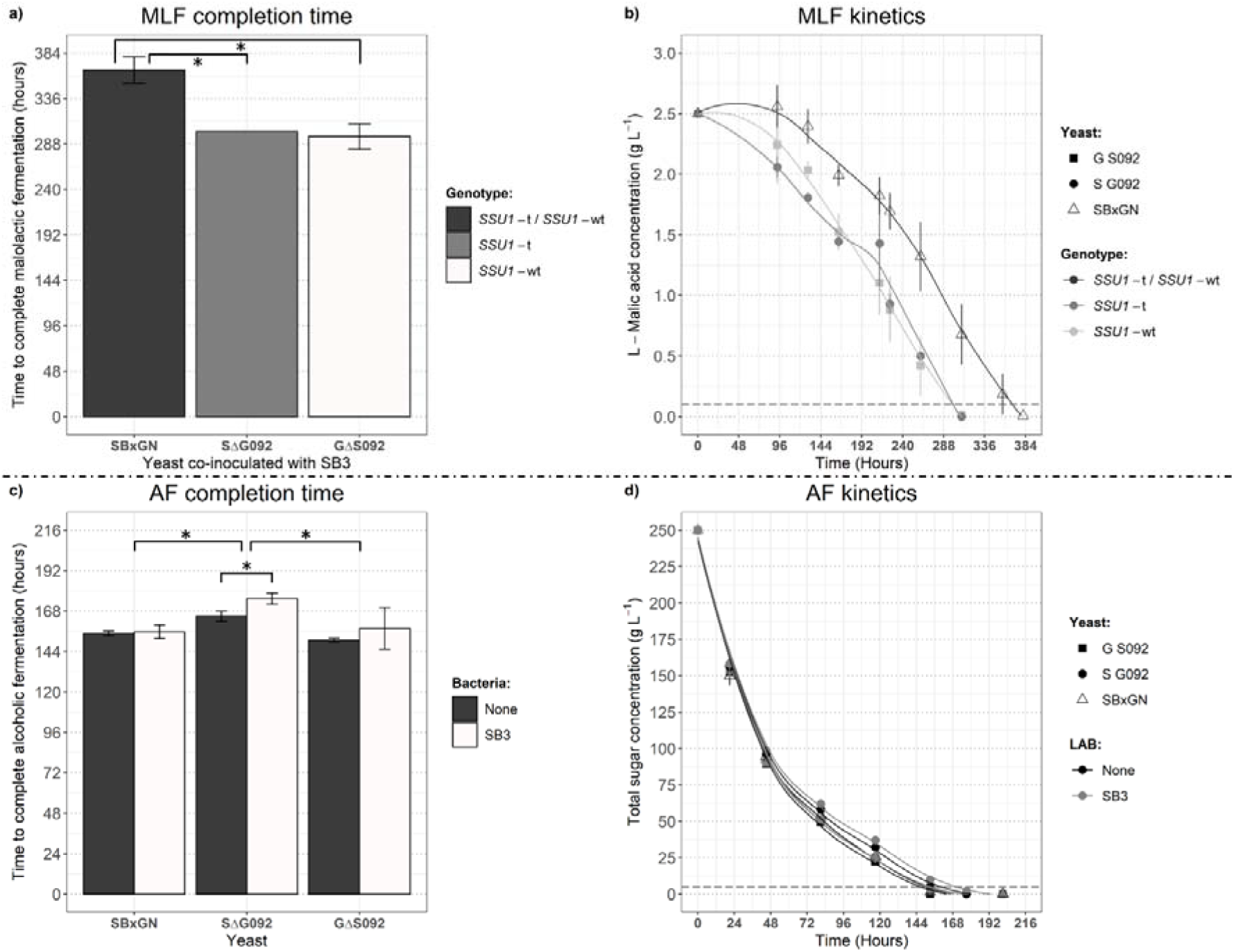
**a)** Time taken for SB3 to complete malolactic fermentation during co-inoculation with SBxGN, SΔG092 and GΔS092. Bar colour indicates the yeast *SSU1* genotype. Significant differences (ANOVA, Tukey post-hoc; *p* < 0.05) between yeast are indicated by *. **b)** L-malic acid consumption over time by SB3 when co-inoculated with SBxGN (triangle), SΔG092 (circle) or GΔS092 (square). Lines demonstrate fitting of local polynomial regression using the R-loess function, span = 0.75. **c)** Time to complete alcoholic fermentation for yeast alone (filled bars) vs yeast co-inoculated with SB3 (colourless bars). Significant differences (ANOVA, Tukey post-hoc; *p* < 0.05) are indicated by *. **d)** Total sugar consumption over time by SBxGN (triangle), SΔG092 (circle) and GΔS092 (square) alone (dark), or co-inoculated with SB3 (light). Lines demonstrate fitting of local polynomial regression using the R-loess function, span = 0.8. Values are the mean of triplicates and error bars are the standard deviation

In addition to MLF, AF kinetics of isogenic hybrids co-inoculated with and without SB3 were also determined. The AF completion time (*tend-AF*) was significantly longer for SΔG092 compared to SBxGN and GΔS092 (Fig. 4c and d), but was unaffected by the presence of LAB. Additionally, in co-inoculated fermentations, SΔG092 (functional GN allele) had a significantly slower fermentation (Fig. 4c; ANOVA, Tukey post-hoc, *p* < 0.05, Table S2: Online resource 2). Differences in AF kinetics between *SSU1* hemizygous hybrids have been previously reported, where the GN allele correlated to reduced lag phase, increased cell viability (Zimmer et al. 2014) and longer fermentation completion time (Peltier et al. 2018b) in connection with the SO_2_ content of grape juice. For the present work, no sulfite was added to the grape juice, removing the influence of this antimicrobial agent. Therefore, the differences of fermentation kinetics observed for the SΔG092 may be in part due to yeast-LAB interactions. This unexpected result suggests that LAB co-inoculation with yeast may impact yeast fermentation fitness and allelic variations of *SSU1* in yeast could modulate such interactions.

## Discussion

### Implementation of QTL mapping for narrowing down genetic regions controlling microbial interactions

Previously, studies investigating yeast-LAB interactions during juice fermentation relied on AF and MLF kinetics and production of volatile and non-volatile compounds (Arnink and Henick-Kling 2005; Comitini and Ciani 2007; Mendoza et al. 2010; Nehme et al. 2008, 2010). Many combinations of yeast and LAB, either sequentially or co-inoculated in juice or wine, have revealed that yeast-LAB compatibility is strain specific (Comitini et al. 2005; Comitini and Ciani 2007; Muñoz et al. 2014). Considering the differences reported for different yeast and LAB strains, and production of different metabolites by yeast strains, there is no question of the influence that yeast genetic makeup has on co-inoculation outcomes. In addition to studies on strain combination and metabolite production, further work has revealed differential *S. cerevisiae* gene expression in response to co-inoculation with *O. oeni* (Rossouw et al. 2012). Though insightful, none of these works have identified specific genetic differences between yeast strains that may influence compatibility with *O. oeni*. Hence, the present work has laid a foundation for understanding how *S. cerevisiae* genetic makeup can impact MLF outcomes during co-inoculation with *O. oeni*. The application of a QTL mapping strategy was used to find specific influences of yeast genetic makeup on yeast-LAB interactions.

In the past, QTL mapping has been used for investigating plant-pathogen interactions unveiling links between plant genotype and pathogen resistance (Chen et al. 2010; Decroocq et al. 2005; Eun et al. 2016). In the current work, the same strategy was applied to the two main species involved in winemaking: *S. cerevisiae* and *O. oeni*, to delineate the influence of yeast genetic background in microbial interactions.

QTL mapping has been used numerous times for *S. cerevisiae* to determine the genotypic traits that influence yeast AF completion (Marullo et al. 2019), acetic acid production (Marullo et al. 2007b; Salinas et al. 2012), nitrogen uptake (Brice et al. 2014; Jara et al. 2014) and aroma compound production (Eder et al. 2018; Huang et al. 2014; Roncoroni et al. 2011; Steyer et al. 2012) which are important for fermentation progress and wine quality. However, to the best of our knowledge, QTL mapping has not been used to study yeast-bacteria interactions in wine. To achieve this task, we used the BN progeny clones (pop BN, Table 1) that have been previously used for investigating the genetic determinism of fermentation traits (Martí-Raga et al. 2017; Zimmer et al. 2014). Each progeny clone was co-inoculated (i.e. exposed) to LAB during fermentation and the percentage of L-malic acid consumed was measured. For the *S. cerevisiae* strains alone, L-malic acid at the end of fermentation ranged from 44% consumption to 10% production when compared to the starting concentration of 2.5 g L^-1^ (Fig. 1b; Table S1: Online resource 1) and the ability of yeast to consume or produce this amount of L-malic acid is in agreement with previous findings (Delcourt et al. 1995; Peltier et al. 2018a; Yéramian et al. 2007). The continuous distribution of L-malic acid consumption or production observed among the yeast progeny suggests that this trait is controlled by many genes. A study detailing those genes is currently under preparation (Peltier, pers. comm.).

Since *S. cerevisiae* strains can consume some of the L-malic acid present in grape must, the specific consumption of L-malic acid by LAB was determined by subtracting L-malic acid concentration in yeast-alone fermentations from values in co-inoculations. This computed trait (*Malic_acid_LAB_consumed*) was used to estimate the influence of yeast strain on *O. oeni* strain SB3’s ability to complete MLF. The ability of SB3 to complete MLF in co-inoculated conditions showed a gaussian distribution, suggesting that several *S. cerevisiae* genes control this phenotype (Fig. 2a). Although the number of progenies tested was low for achieving QTL mapping (67 clones), we implemented a linkage analysis that revealed a major QTL (XVI_374) with a LOD score above an FDR threshold of 5%. The marker for this major QTL was located within *SSU1*, giving reason to perform follow-up work with *SSU1* hemizygote strains. Beside this major QTL, one other loci (XV_162) reached a lower FDR threshold of approximately 10%. The QTL XV_162 marker corresponds to *PHM7*, a gene of unknown function. However, this marker was physically linked with the major QTL XVI_374 due to the presence of the XV-t-XVI translocation that was previously identified in this genetic background (Zimmer *et al*. 2014). Therefore, our strategy was useful for narrowing down two unlinked genetic regions that impacted the success of MLF by SB3 in co-inoculated fermentations.

### Putative role of *SSU1* gene in yeast-LAB interactions

Once identified by linkage analysis, QTLs can be dissected at the gene and single-nucleotide polymorphism (SNP) level to further understand the underlying molecular genetic mechanism of phenotypic traits (Peltier et al. 2019). However, this functional characterisation is a complex task that cannot always be achieved. In the present study, there was no clear demonstration of the impact of natural allelic variations linked to SB3 MLF completion in co-inoculated fermentations.

The QTLs identified in this study link two markers, XVI_374 and XV_162, to the phenotype investigated. These two QTLs belonging to chromosomes XV and XVI must be considered as a single locus since they are physically linked by a reciprocal translocation event, XV-t-XVI, in the parental strain GN. This translocation is detected in numerous wine related strains (Marullo et al. 2020; Treu et al. 2014; Zimmer et al. 2014) and can impact numerous yeast phenotypes including lag phase duration, cell viability (Zimmer et al. 2014), growth rate in the presence of SO_2_ (Marullo et al. 2020) and alcoholic fermentation kinetics (Peltier et al. 2018b). These phenotypes have been attributed to the modification of the promoter sequence of the gene *SSU1*, that encodes Ssu1p, an intermembrane transporter responsible for *S. cerevisiae* sulfite efflux (Park and Bakalinsky 2000). In GN, the translocation results in decreased distance between the *ADH1* promoter region and *SSU1*, triggering increased and constitutive *SSU1* expression (Zimmer et al. 2014). Interestingly, two other gross chromosomal rearrangements related to the *SSU1* promoter have been described in the past (García-Ríos et al. 2019; Pérez-Ortín et al. 2002) and were associated with increased *SSU1* expression as well as increased sulfite resistance (García-Ríos et al. 2019; Marullo et al. 2020). These chromosomal rearragements have facilitated adaptation of wine yeast, enabling increased SO_2_ efflux from the yeast cell in oenological conditions, and has been reviewed recently (Divol et al. 2012; García-Ríos and Guillamón 2019). In terms of co-inoculation, yeasts that efficiently export sulfite could negatively impact *O. oeni* growth and MLF progress, as SO_2_ can inhibit the internal ATPase of this lactic acid bacterium (Carrete et al. 2002). This is supported by the linkage analysis since the GN allele had a deleterious effect on MLF completion (Fig. 2d). We tested this hypothesis by reciprocal hemizygosity assay, including fermentations with and without LAB co-inoculation and determining the percentage of L-malic acid consumed by LAB. Although MLF was completed in all the co-inoculated fermentations, a strong haploinsufficiency effect was observed between the native hybrid SBxGN and the two hemizygous hybrids (Fig. 4a). Thus, the lack of one copy of *SSU1* within yeast can enable considerably reduced duration of MLF by LAB, no matter which *SSU1* allele is deleted (Fig. 4a and b). This result suggests that the *SSU1* gene may play a role in the phenotype investigated. However, the comparison of hemizygous hybrids fails to confirm the inhibitor role of the *SSU1* GN allele specifically, compared to SB. This negative result may be due to a masking effect of *SSU1* haploinsufficiency, suggesting that the negative impact of the GN allele may require two functional *SSU1* alleles to be fully expressed.

Intriguingly, the AF kinetics of the hemizygous strain SΔG092, which carries the *SSU1* GN allele, was slightly delayed compared to GΔS092 and SBxGN (Fig. 4c and d). In addition, when SΔG092 was co-inoculated with LAB, this delay was longer (up to 20 hours) suggesting a possible inhibition of yeast AF by bacteria. This deleterious effect of the GN allele on fermentation kinetics has been previously reported (Peltier et al. 2019) and was attributed to the increase of external sulfite in the fermented medium, a direct effect of increased SO_2_ efflux. However, in our conditions, sulfite concentration was below the detection threshold, and no SO_2_ was added to the grape juice.

The inability to confirm the QTL using the reciprocal hemizygosity assay could be due to *SSU1* not being the main gene involved in MLF completion by LAB or AF activity. In fact, other genes genetically linked to the GN allele of *SSU1* could be involved in those phenotypes. Among them, *ADH1* may be an interesting candidate. *ADH1* encodes an alcohol dehydrogenase that is located on chromosome XV, but is also physically associated with the *SSU1* gene in the XV-t-XV translocation genotype. In the first hours after yeast inoculation, expression of *ADH1* is not fully active. Ethanol production is delayed and replaced by glycerol-pyruvic fermentation in order to cope with the depletion of NAD^+^ in the cytosol (Ribereau-Gayon et al. 2006). We hypothesise that the XV-t-XVI translocation could also modify the expression of *ADH1*, at least in the early stage of yeast culture. Decreased *ADH1* expression in the early phase of AF could lower alcohol dehydrogenase activity, increasing the accumulation of acetaldehyde. This phenomenon has been previously observed for different wine starters including VL1, the parental strain of GN (Cheriati et al. 2010). The biological implication of acetaldehyde utilisation by wine LAB in free or bound form is not yet fully understood, apart from the liberation of free SO_2_ from the acetaldehyde-SO_2_ complex (Liu and Pilone 2000). High levels (> 100 mg L^-1^) of acetaldehyde may inhibit growth of LAB (El-Gendy et al. 1983). Thus, the impact of the XVI_374 QTL observed in the SBxGN progeny might be due to other genetic polymorphisms related to the XV-t-XVI translocation, such as the modification of the *ADH1* promoter region. In future, additional experimental efforts will be required to further investigate these molecular causes.

In conclusion, for the first time, yeast genetic background was assessed for its role in yeast-LAB compatibility during fermentation. The impact of *SSU1* haploinsufficiency on LAB ability to complete MLF was clear, but further work is needed to understand the role of the XV-t-XVI translocation on MLF outcomes. The inclusion of more progeny and use of a different genetic background may help identify other genes that result in phenotypic differences in yeast-LAB co-inoculations. Further work may also reveal the underlying cause of the major QTL identified in this study, which was not able to be confirmed by the *SSU1* hemizygote strains. However, the influence of *SSU1* in this work does add to the understanding of the pleiotropic role of *SSU1*, since it has been reported to impact yeast AF, growth and SO_2_ production, and now also has the potential to impact co-inoculation outcomes with LAB. This work begins to unravel the complexity of *S. cerevisiae* genetic differences that can lead to a phenotype that impacts *O. oeni* during co-inoculation. Understanding the delicate interplay between genotype and phenotype can create opportunities for wine yeast manufacturers to develop yeast that work effectively with LAB, without negatively impacting yeast AF performance.

## Supporting information

Online resource 1

Online resource 2

## Declarations

### Funding

This work was supported by Australia’s grape growers and winemakers through their investment body, Wine Australia, with matching funds from the Australian Government. LB was supported by joint scholarships from The University of Adelaide and Wine Australia (AGW Ph 1510). JS was supported by Wine Australia project funding (UA1707). JS, KS and VJ are supported by The Australian Research Council Training Centre for Innovative Wine Production (www.ARCwinecentre.org.au; project number IC170100008), which is funded by the Australian Government with additional support from Wine Australia and industry partners. The University of Adelaide is a member of the Wine Innovation Cluster in Adelaide (http://www.thewaite.org/waite-partners/wine-innovation-cluster/). PM and EP are supported by Biolaffort (Laffort® Research & Development subsidiary) for this project, in addition PM received a grant from Aquitaine Region (Sesam Project) for genome sequencing and QTL analysis.

### Ethical approval

This article does not contain any studies with human participants or animals performed by any of the authors.

### Availability of data

Raw data may be supplied upon request, at the discretion of the corresponding authors.

### Authors’ contributions

All authors contributed to the study conception and design. LB carried out laboratory fermentation experiments; PM and EP provided yeast strains. Data analysis and interpretation was performed by LB, EP and PM. LB wrote the manuscript and EP, JS, KS, JGM, VJ and PM reviewed and revised the manuscript. All authors read and approved the final manuscript.

## Acknowledgements

We thank Hélène Mesnage for her assistance in performing the QTL fermentation experiment. We also acknowledge Nick van Holst Pellekaan for capturing images of the automated fermentation platform.

## Conflicts of interest

EP and PM are employed by Biolaffort. This does not alter the authors’ adherence to all the journal policies on sharing data and materials.

## Notes

### Competing Interest Statement

The authors have declared no competing interest.

