## Supplementary material for "QTL mapping: an innovative method for investigating the genetic determinism of yeast-bacteria interactions in wine": Online resource 2

bioRxiv

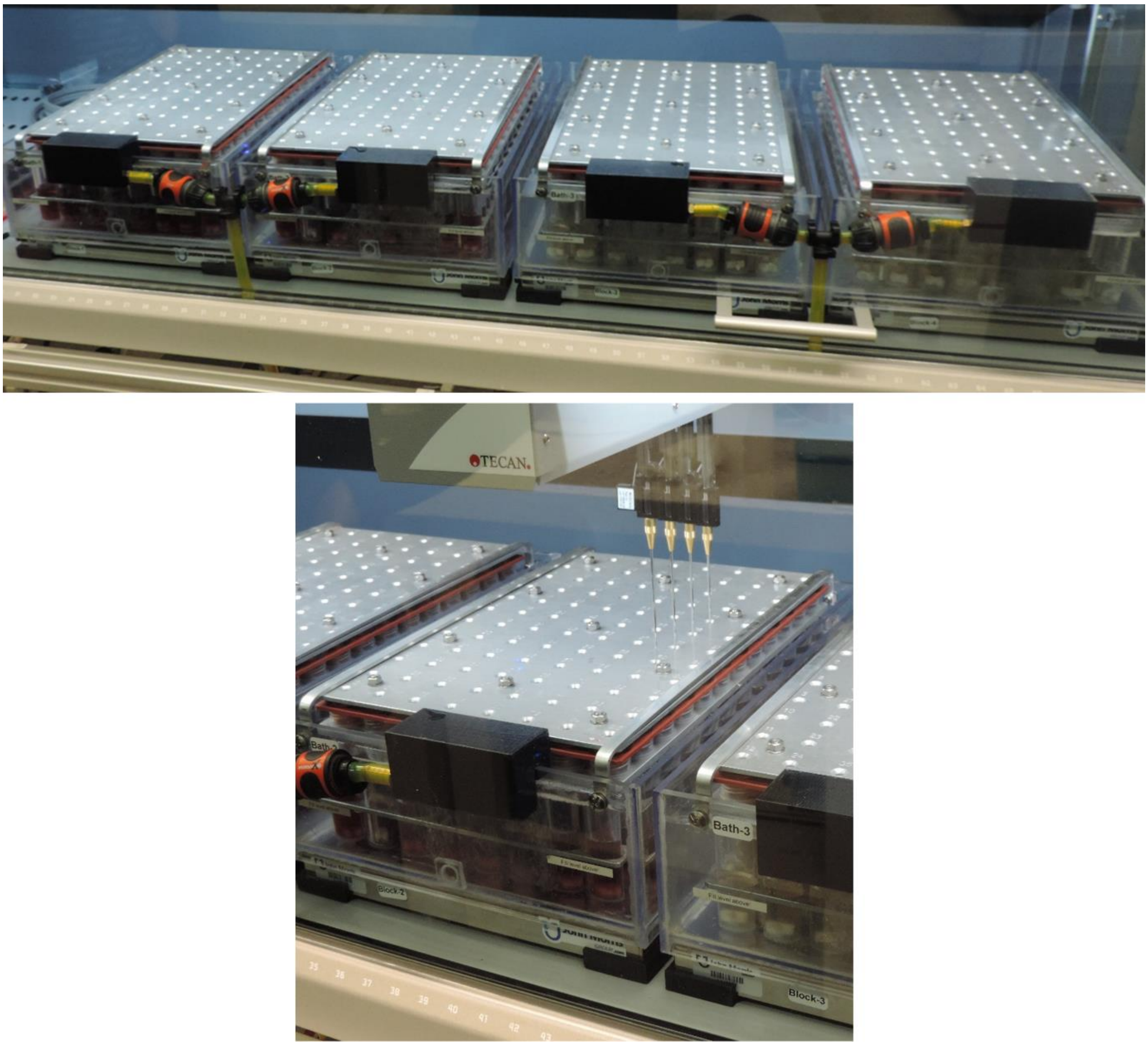


**Fig. S1** 384 automated fermentation platform. Set-up included 4 x 96 tube blocks, each temperature controlled by water baths and individual tube mixing by magnetic stir bars (top). Fermentations were sampled aseptically via septum-sealed port using an automated overhead needle system (bottom)

**Table S2**: ANOVA results for SBxGN and hemizygote fermentations. *P*-value and variance are displayed for the individual tests for SSU1 allele (SB, GN or both: SB/GN), presence of bacteria (co-inoculation with SB3 or no co-inoculation), and combinatorial effect of allele and bacteria. Post-hoc analysis was used to determine groups (designated by the letters) for allele and bacteria variables. Measures tested were tend-AF, t35-AF and s50-80-AF which represent time to complete AF, time to complete 35% of AF and slope value for points between t50-AF and t80-AF, respectively.

| | **SBxGN** | **tend-AF** | **t35-AF** | **s50-80-AF** |
|---|---|---|---|---|
| ***p*-value** | *SSU1* allele | 0.001 | 0.432 | 0 |
| | Bacteria (co-inoculated or not) | 0.042 | 0.047 | 0.254 |
| | Combinatorial effect of allele and bacteria | 0.383 | 0.906 | 0.14 |
| **Variance observed** | *SSU1* allele | 61 | 10 | 77 |
| | Bacteria (co-inoculated or not) | 11 | 26 | 2 |
| | Combinatorial effect of allele and bacteria | 4 | 1 | 6 |
| **Post-hoc group: *SSU1* allele** | GN | a | a | a |
| | SB/GN | b | a | b |
| | SB | b | a | b |
| **Post-hoc group: Bacteria (co-inoculated or not)** | No co-inoculation | b | b | a |
| | Co-inoculation with SB3 | a | a | a |
